# Exploring SNP Filtering Strategies: The Influence of Strict vs Soft Core

**DOI:** 10.1101/2024.08.26.609800

**Authors:** Mona L. Taouk, Leo A. Featherstone, George Taiaroa, Torsten Seemann, Danielle J. Ingle, Timothy P. Stinear, Ryan R. Wick

**Affiliations:** Department of Infectious Diseases, The University of Melbourne at the Peter Doherty Institute for Infection and Immunity, Melbourne, Victoria, Australia; Department of Microbiology and Immunology, The University of Melbourne at the Peter Doherty Institute for Infection and Immunity, Melbourne, Victoria, Australia; Macroevolution and Macroecology Group, Research, School of Biology, Australian National University, Canberra, ACT, Australia; Centre for Pathogen Genomics, The University of Melbourne, Melbourne, Victoria, Australia

## Abstract

Phylogenetic analyses are crucial for understanding microbial evolution and infectious disease transmission. Bacterial phylogenies are often inferred from single nucleotide polymorphism (SNP) alignments, with SNPs as the fundamental signal within these data. SNP alignments can be reduced to a ‘strict core’ by removing those sites which do not have data present in every sample. However, as sample size and genome diversity increase, a strict core can shrink markedly, discarding potentially informative data. Here, we propose and provide evidence to support the use of a ‘soft core’ that tolerates some missing data, preserving more information for phylogenetic analysis. Using large datasets of *Neisseria gonorrhoeae* and *Salmonella enterica* serovar Typhi, we assess different core thresholds. Our results show that strict cores can drastically reduce informative sites compared to soft cores. In a 10,000-genome alignment of *Salmonella enterica* serovar Typhi, a 95% soft core yielded 10 times more informative sites than a 100% strict core. Similar patterns were observed in *Neisseria gonorrhoeae*. We further evaluated the accuracy of phylogenies built from strict and soft-core alignments using datasets with strong temporal signals. Soft-core alignments generally outperformed strict cores in producing trees displaying clock-like behaviour; for instance, the *Neisseria gonorrhoeae* 95% soft core phylogeny had a root-to-tip regression *R^2^* of 0.50 compared to 0.21 for the strict-core phylogeny. This study suggests that soft-core strategies are preferable for large, diverse microbial datasets. To facilitate this, we developed *Core-SNP-filter* (github.com/rrwick/Core-SNP-filter), an open-source software tool for generating soft-core alignments from whole-genome alignments based on user-defined thresholds.

**IMPACT STATEMENT:** This study addresses a major limitation in modern bacterial genomics – the significant data loss observed in large datasets for phylogenetic analyses, often due to strict-core SNP alignment approaches. As bacterial genome sequence datasets grow and diversity increases, a strict-core approach can greatly reduce the number of informative sites, compromising phylogenetic resolution. Our research highlights the advantages of soft-core alignment methods which tolerate some missing data and retain more genetic information. To streamline the processing of alignments, we developed *Core-SNP-filter* (github.com/rrwick/Core-SNP-filter), a publicly available resource-efficient tool that filters alignments to informative and core sites.

**DATA SUMMARY:** All genomic sequence reads used in this study were already publicly available and accessions can be found in Supplementary Dataset 1. Supplementary methods and all code can be found in the accompanying GitHub repository: (github.com/mtaouk/Core-SNP-filter-methods).

## INTRODUCTION

Appropriately resolving the phylogenetic relationships between organisms is a fundamental aspect of many scientific disciplines, including ecology, microbiology and evolutionary biology (1). Defining phylogenetic relationships using whole-genome sequencing (WGS) serves as a cornerstone application in clinical and public health settings for tracking the spread of pathogens and pinpointing outbreaks (2–7).

Conventionally, phylogenetic studies using WGS rely on reference-based techniques. In this approach, single nucleotide polymorphisms (SNPs) are identified for each sample by aligning sequencing reads against a reference genome and calling variants using bioinformatic tools such as *Snippy* (github.com/tseemann/snippy). These results are then combined to generate a whole-genome pseudoalignment (‘alignment’ for brevity) encompassing all relevant samples. If appropriate, recombination events and problematic areas of the genome can be masked from the alignment using tools such as *Gubbins* or *ClonalFrameML* (8, 9). This alignment is then traditionally refined to produce what is known as a ‘strict core’, which includes only the SNP sites present in 100% of samples (see github.com/sanger-pathogens/snp-sites as an example). This strict-core approach is commonly used to generate the data required for phylogenetic inference, calculating SNP distances to infer the relatedness of genomes and patterns of transmission, and being the foundation for temporal phylogenetic inferences, especially in outbreak investigations and evolutionary studies (10–17).

The rapid decrease in sequencing costs has revolutionised our capacity for large-scale whole-genome analyses (18). Contemporary bacterial studies now routinely involve WGS for thousands of samples, and public datasets number in the hundreds of thousands for some species. This necessitates revisiting methods for large-scale analyses, both within and across species. This is evidenced by studies employing methods other than whole-genome alignments to determine the genetic relatedness of isolates (19–22), with questions arising regarding the informativeness of alignments and the challenges of dealing with increasingly large datasets (23). As more genomes are included, the strict-core alignment, consisting of sites present in all genomes, can only diminish in size due to the increasing likelihood of missing data in at least one genome, thereby reducing the overall resolution. This highlights the long-standing debate on the impact of excluding potentially informative sites (24, 25). Some contemporary studies have addressed this concept by employing a ‘soft core’ alignment by filtering the whole-genome alignment to include SNP sites present in fewer than all samples (<100%), although these approaches are not routine in bacterial genomics (6, 26, 27).

In this study, we evaluate the effectiveness of strict and soft-core approaches for filtering whole-genome alignments and phylogenetic inferences using four priority bacterial species. To do so, we introduce *Core-SNP-filter* (github.com/rrwick/Core-SNP-filter), a tailored software tool which allows for efficient manipulation of large SNP alignments. Through these analyses, we aim to illuminate best practices for researchers seeking to accurately resolve phylogenetic relationships in bacteria.

## METHODS

### Development and implementation of Core-SNP-filter

To allow for the efficient filtering of core SNPs to user-defined thresholds, we developed *Core-SNP-filter*, which takes a FASTA alignment as input and outputs a reduced FASTA alignment based on core filtering settings. *Core-SNP-filter* accepts any FASTA alignment as input, including whole-genome alignments (such as output by *Snippy* or *SKA2*), gene-based alignments (such as output by *Roary* or *Panaroo*), and alignments with or without invariant sites. Additionally, we note that alignments without invariant sites and no missing or ambiguous data will remain unchanged by the filtering process.

*Core-SNP-filter* can discard invariant sites (controlled by the –e option) and non-core sites (controlled by the –c option). A site is determined to be invariant if the number of unique unambiguous bases (A, C, G or T) in that site is zero or one. A site is determined as core if the fraction of bases in that site which are unambiguous exceeds the user-supplied value (e.g. –c 0.95). All characters other than the unambiguous bases (e.g. N, –, X, etc.) are treated the same (see Fig. 1 for an illustrated example).

**Figure 1:**
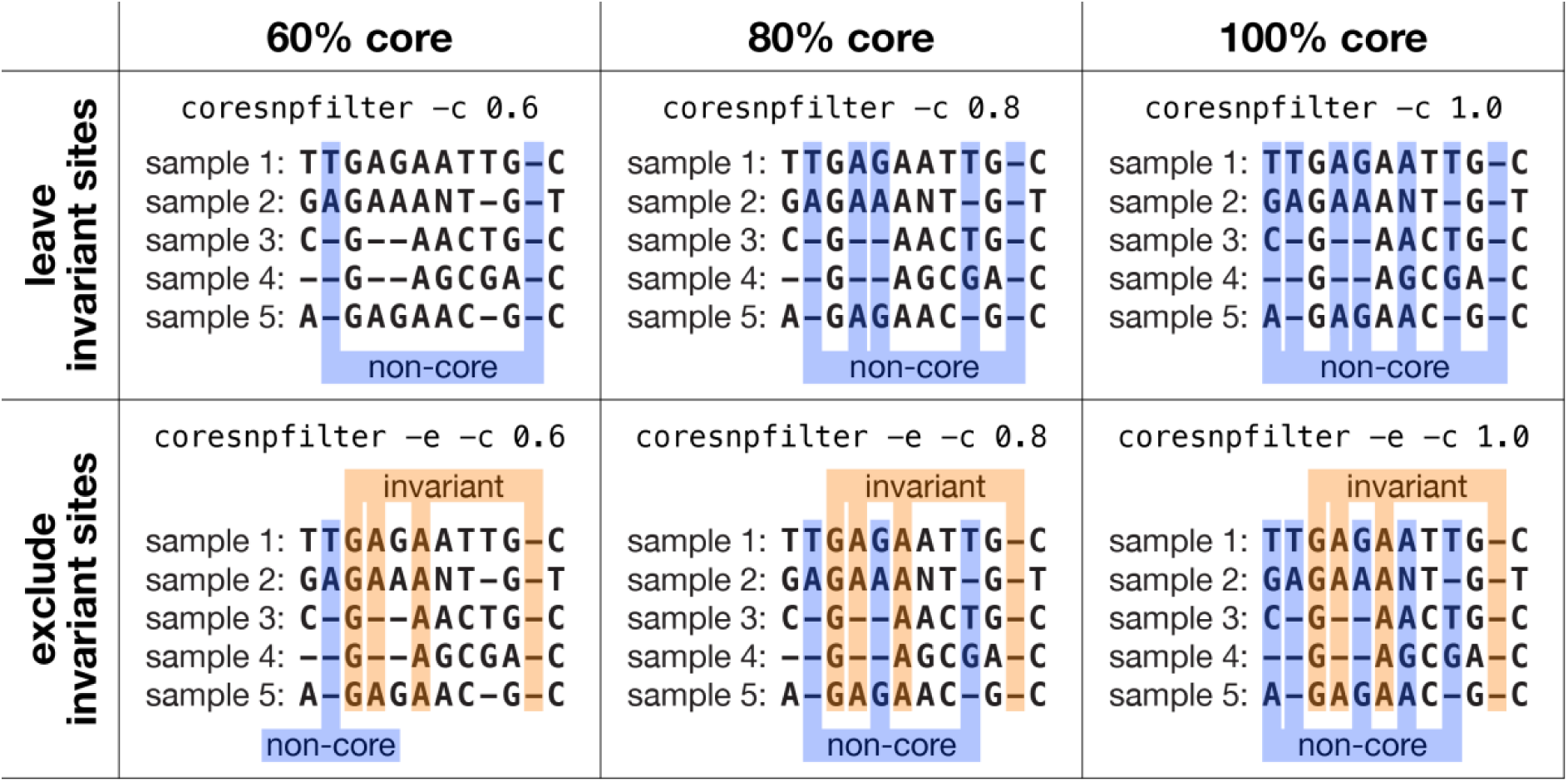
Schematic of the core filtering process at various thresholds. A visual depiction of how Core-SNP-filter processes sites with different settings. The highlighted sites are the ones that Core-SNP-filter will remove, i.e. the output will consist only of the non-highlighted sites. Core-SNP-filter uses ≥ for core site assessment, e.g. with –c 0.8, a site with 4/5 unambiguous bases is considered core. All non-A/C/G/T characters (e.g. N and –) are treated equally. A site can be both invariant and non-core, and when both –e and –c are used, invariant sites are filtered first, so some non-core sites may shift to the invariant category.

*Core-SNP-filter* is implemented in Rust and is resource-efficient. It is available on GitHub (github.com/rrwick/Core-SNP-filter) with thorough usage instructions and a sample dataset.

### Large-scale genome analysis using PathogenWatch data

For two priority pathogens, *Neisseria gonorrhoeae* (n = 11,129) and *Salmonella enterica* serovar Typhi (n = 10,991), we downloaded the reads for all genomes available on PathogenWatch, an online database for processing and visualising bacterial genome sequences in phylogenetic and geographical contexts with associated accessions and published studies (details in Supplementary Dataset) (28). In addition to being highly represented on PathogenWatch, these two species were selected because *N. gonorrhoeae* is naturally competent for genetic transformation and therefore represents a highly recombinogenic and diverse species, while *S.* Typhi has lower genetic diversity and less recombination (29, 30). *Snippy* v4.6.0 was used with default settings to call variants, using *N. gonorrhoeae* NCCP11945 (accession: CP001050.1) and *S.* Typhi str. CT18 (accession: NC_003198.1) as reference genomes (31, 32). Genomes with more than 10% ambiguous bases (N characters) in their *Snippy* output sequence were excluded from downstream analyses (see Supplementary Methods).

For both species, we generated a series of alignments with progressively larger genome counts by incrementally adding genomes. Starting with an alignment of 25 genomes, we added genomes in increments of 25, creating alignments of 50, 75, and 100 genomes, and then continued with increments of 250 genomes, up to 10,000 genomes. Each alignment includes all genomes from the previous alignment, ensuring consistency across alignments and resulting in a total of 44 datasets with varying genome counts. For each of these 44 alignments, a core alignment was produced at thresholds of 50%, 60%, 70%, 80%, 90%, 95%, 96%, 97%, 98%, 99% and 100% using *Core-SNP-filter*, once removing invariant sites and a second time keeping all invariant sites. From each, the alignment length (number of remaining sites) was calculated. This entire process was repeated ten times for each species, with a new random seed used for each repetition, and the reported results are the mean of all ten replicates. Additionally, for one set of *S.* Typhi alignments ranging from 25 to 2000 genomes, *Gubbins* v2.4.1 was used to identify regions of recombination (8). For each alignment *Core-SNP-filter* was used on the polymorphic sites identified by *Gubbins to* produce core alignments at thresholds of 50%, 60%, 70%, 80%, 90%, 95%, 96%, 97%, 98%, 99% and 100%.

To investigate the effect of changing between strict and soft-core thresholds on related alignments, core SNP alignments were generated for 22 individual *N. gonorrhoeae* and 18 *S.* Typhi studies at both 100% and 95% thresholds (7, 16, 17, 33–69). Genomic reads were downloaded from PathogenWatch corresponding to the published studies (Supplementary Methods). For each study, variants were called using *Snippy* with the *N. gonorrhoeae* NCCP11945 reference genome or *S. Typhi* str CT18 reference genome, a pseudoalignment was made with *Snippy-core*, and *Core-SNP-filter* was used to make 100% strict-core and 95% soft-core alignments. The number of variant sites in each core alignment was then counted and compared between the two thresholds. For each study alignment, the pairwise SNP distances between genomes were calculated with *snp-dists* v0.8.2 (github.com/tseemann/snp-dists).

### Evaluation of core thresholds for phylodynamics with public datasets

Four publicly available datasets were used to test core thresholds on a temporal regression analysis: 154 *Streptococcus pneumoniae* genomes (70), 192 *N. gonorrhoeae* genomes (17), 88 *Staphylococcus aureus* genomes (10) and 96 *Salmonella enterica* serovar Kentucky genomes (13). These datasets were chosen because they each span a range of dates and contain a strong temporal signal, as demonstrated in their original publications.

For each dataset, we downloaded all available read sets and the reference genome used in the dataset’s publication (see Supplementary Methods). Quality checks were performed on the reads from the *S. pneumoniae* dataset (excluding samples unable to assemble well, see Supplementary Methods), as these reads were from 2011 and lower quality than the other three datasets. *Snippy* v4.6.0 was used with default settings to produce whole-genome pseudoalignments using the reference genome. *Snippy-core* was used to merge these into a combined alignment for each study. *Gubbins* v3.2.1 was used to generate a pseudoalignment with putative recombinant regions masked (8). We then used *Core-SNP-filter* to produce an invariant-free alignment at each possible core threshold (e.g. 0/n, 1/n, 2/n, … (n-1)/n, n/n) for both the pre-*Gubbins* and post-*Gubbins* alignment.

We then used *IQ-TREE* v2.0.3 to produce three independent trees from each pseudoalignment using the TVM+F+ASC+R2 substitution model, the best-supported model identified by *IQ-TREE’s* inbuilt ModelFinder throughout these datasets (71). Each tree was rooted by selecting the root position maximising *R^2^* for root-to-tip regression using *ape* v5.7.1 (72). For each pseudoalignment/tree, we extracted the number of variant sites and *R*² of the root-to-tip regression (a measure of the temporal signal strength for a strict molecular clock). We expect higher values of *R*² at the 95% core threshold compared to the 100% threshold because reduced polytomy should lead to better fit in root-to-tip regression by informing the spread of root-to-tip distances on the y-axis. Robinson-Foulds (RF) distances were calculated using *phangorn* v2.12.1 to compare the topologies of phylogenies generated with a 100% strict-core threshold to phylogenies generated at each possible core threshold across each of the four datasets with three replicates (73, 74). RF distances were normalised such that a value of one represents maximally dissimilar trees, while a value of zero represents identical topology.

### Benchmarking of Core-SNP-filter against similar tools

To evaluate the computational performance of *Core-SNP-filter*, we conducted benchmarking tests relative to three other tools: *Goalign* v0.3.7, *SNP-sites* v2.5.1 and *TrimAl* v1.4.1 (75–77). We quantified the time and RAM required for two tasks: converting full alignments to no-invariant-sites alignments and converting no-invariant-sites alignments to 95% core alignments. *Core-SNP-filter* and *Goalign* can perform both tasks, while SNP-sites can only perform the first task and *TrimAl* can only perform the second. Each tool was cloned from GitHub and built from source code on a system equipped with an AMD EPYC 7742 CPU and 512 GB of RAM. The test dataset consisted of a 10,000-genome *S.* Typhi pseudoalignment, which was 4,809,037 base pairs in length and 47 GB in size. The sequences contained the characters –, A, C, G, T, N, a, c, g, t and n. Genome counts ranging from 10 to 10,000 were tested, spaced evenly on a logarithmic scale.

The benchmarking process involved subsampling the initial alignment to create datasets of varying genome counts. For each dataset, we used the tools to remove invariant sites and then to produce a 95% core alignment. Each test was repeated eight times, and the minimum time and RAM usage were recorded. MD5 checksums of the output files ensured consistency of results across different tools.

### Statistical analyses and data visualisation

All statistical analyses were performed in *RStudio* v2022.12.0+353 using *R* v4.2.2. All figures were made using *ggplot2* v3.4.0.

## RESULTS

### Revisiting the appropriateness of strict-core filtering for large and diverse datasets

We evaluated the impact of core thresholds on alignment size and number of informative sites for phylogenetic analyses for two species with more than ten thousand publicly available genomes: *N. gonorrhoeae* and *S*. Typhi. For *S.* Typhi, we analysed 10,991 genomes collected between 1935 and 2020 from 95 countries, forming a geographically and temporally diverse dataset (accessions in Supplementary Dataset 1). These genomes were sequentially subset into 44 alignments varying by number of genomes (see Methods for details). Similarly, for *N. gonorrhoeae*, 11,129 genomes collected between 1928 and 2018 from 62 countries were also subset into 44 alignments varying by number of genomes (accessions in Supplementary Dataset 1).

For both *N. gonorrhoeae* and *S.* Typhi, the 100% strict-core resulted in progressively fewer alignment sites as the number of genomes increased (Fig. 2a-b, Supplementary Fig. 1, Supplementary Dataset 2). At all other core thresholds tested (50–99%), the number of sites remained stable as the number of genomes increased. For example, an alignment of 10,000 genomes at a strict threshold of 100% resulted in a mean of 256,124 and 633,715 sites for *N. gonorrhoeae* and *S.* Typhi, respectively, compared to a soft threshold of 95%, which resulted in a mean of 1,900,987 and 4,634,869 sites in *N. gonorrhoeae* and *S.* Typhi, respectively.

**Figure 2:**
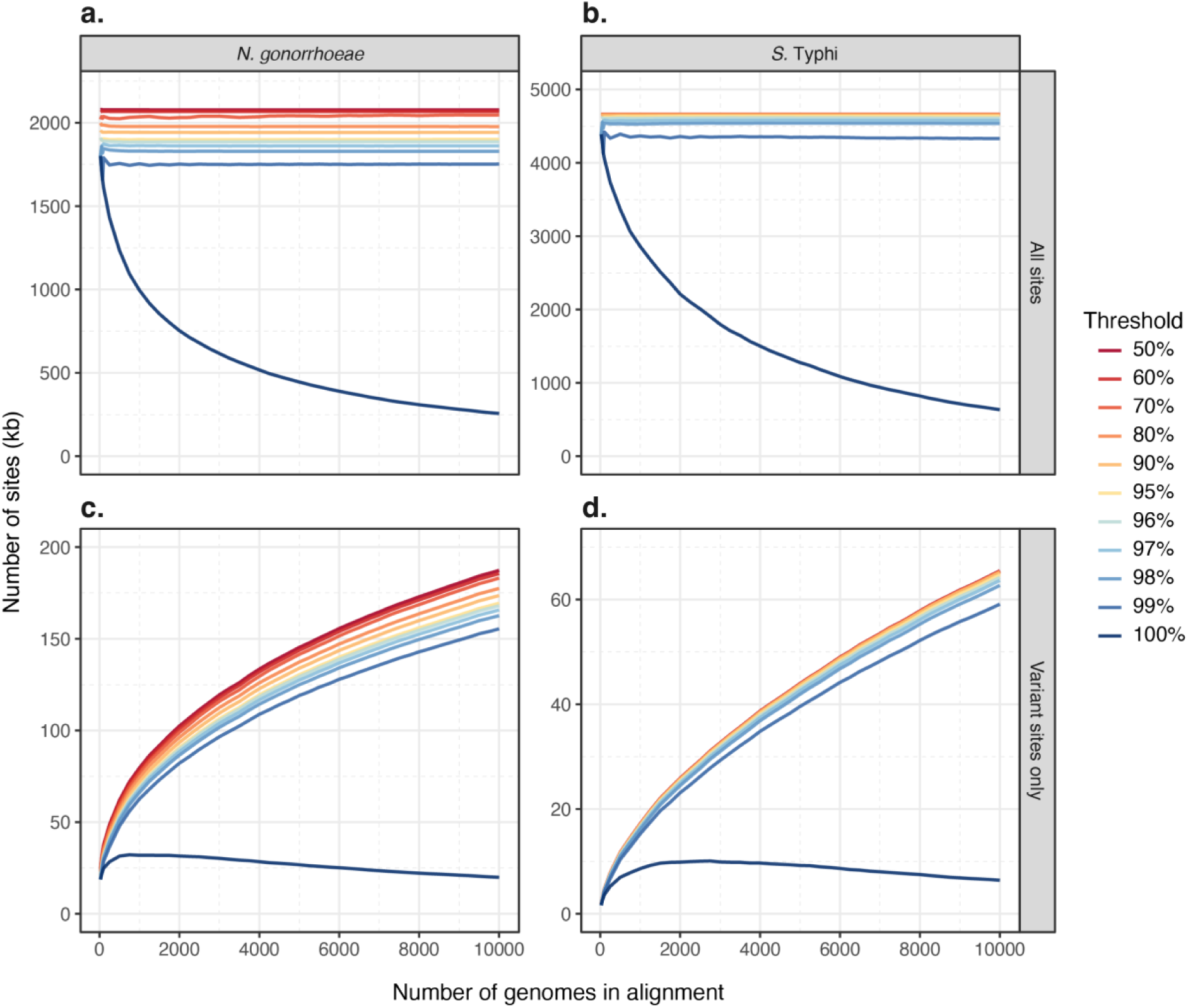
Evaluation of core thresholds on alignment size and variant sites. The number of alignment sites (in kilobases) is plotted against the number of genomes included in each alignment (ranging from 25 to 10,000). Each alignment was processed at core thresholds ranging from 50% to 100% (indicated by line colour). (**a**) All sites in *Neisseria gonorrhoeae* alignments, (**b**) all sites in *Salmonella enterica* subspecies Typhi alignments, (**c**) variant sites only in *Neisseria gonorrhoeae* alignments, and (**d**) variant sites only in *Salmonella enterica* subspecies Typhi alignments. The number of sites was similar across different core thresholds, except for the 100% threshold, which resulted in far fewer sites, particularly at higher genome counts.

A similar trend was observed when considering variant sites. For both *N. gonorrhoeae* and *S.* Typhi, the 100% strict-core resulted in drastically fewer variant sites compared to other thresholds at all alignment sizes (Fig 2c-d). Except for the 100% strict core, the number of variant sites increased with the number of genomes at all tested thresholds, and the number of variant sites increased as the core threshold decreased. For example, an alignment of 10,000 genomes at a strict threshold of 100% resulted in a mean of 19,926 and 6,415 variant sites output for *N. gonorrhoeae* and *S.* Typhi, respectively, compared to a soft threshold of 95%, which resulted in a mean of 169,409 and 64,711 variant sites output in *N. gonorrhoeae* and *S.* Typhi, respectively. Similarly, after removing recombination, the 100% strict-core threshold produced 9,783 variant sites, whereas the 95% soft-core threshold yielded 24,703 variant sites in *S.* Typhi (Supplementary Fig. 2). These findings indicate that considerably more evolutionary signal can be captured using soft-core approaches (<100%) compared to strict-core approaches (100%).

In genomic epidemiology, datasets from individual studies may exhibit reduced diversity and potential biases due to variable temporal, contextual and/or geographic sampling. To assess the potential impacts of varying core thresholds, we collated a set of *N. gonorrhoeae* (n=22) and *S.* Typhi (n = 18) studies and generated alignments for each using both 100% and 95% thresholds (see Methods) (7, 16, 17, 33–69). For all but one study, the number of variant sites output increased in the 95% threshold alignment compared to the 100% threshold alignment (Supplementary Fig. 3-4, Supplementary Table 1-2). In *N. gonorrhoeae*, large and diverse datasets particularly benefited from lowering the core threshold, as shown by Williamson *et al.* 2019 (2,162 genomes), which had the highest increase in variant sites at 511.2% (7). Other studies with substantial increases included Thomas *et al.* 2019 (639 genomes, 107.16%) and Alfsnes *et al.* 2020 (806 genomes, 122.76%) (16, 33). Fifer *et al.* 2018 (97 genomes) and Kwong *et al.* 2018 (75 genomes) exhibited smaller increases of 8.59% and 6.16%, respectively, showing that smaller and/or less diverse datasets may see less benefit from lowering the threshold (40, 42). Ezewudo *et al.* 2015 (18 genomes) showed no change in SNP sites, highlighting that for very small datasets lacking in diversity, threshold adjustments may not impact variant site counts (39). A similar trend was observed in *S.* Typhi, where larger datasets showed more pronounced increases in variant sites at the 95% threshold. For instance, Wong *et al.* 2015 (1,819 genomes) saw a large increase of 216.0% in variant sites, while Da Silva *et al.* 2022 (3,398 genomes) increased by 79.6% (56, 69). In contrast, smaller datasets like Guevara *et al.* 2021 (76 genomes) showed a modest increase from 1,328 to 1,886 variant sites, and Rasheed *et al.* 2020 (26 genomes) had minimal change from 19 to 20 sites (58, 66). This cross-species analysis reinforces that lowering the core threshold can markedly increase variant site counts, especially in large, diverse datasets.

In addition to the observed differences in variant site counts, the 100% core threshold alignment resulted in a higher percentage of identical genome pairs (zero SNP distance) compared to the 95% core threshold alignment across all studies, except for three studies with no identical genomes in either alignment (Supplementary Fig. 5-6, Supplementary Table 1-2). For example, in the Williamson *et al.* dataset (*N. gonorrhoeae*), the proportion of identical genome pairs increased from 0.07% in the 95% core alignment to 2.43% in the 100% core alignment (7). Similarly, in the Wong *et al.* dataset (*S.* Typhi), the percentage rose from 0.06% to 3.82% under the same comparison (69). This trend suggests that the strict-core alignments reduce genomic resolution, even though ambiguous ‘N’ bases are excluded from SNP counts.

### Appropriateness of using a strict-core threshold for small datasets with strong temporal signal

To investigate the effect of varying core thresholds in situations where removing certain sites might be beneficial—such as those inconsistent with the clonal frame of evolution or regions affected by recombination—we analysed four existing datasets. These datasets, characterised by a strong temporal signal and limited geographic diversity, provide an ideal context for assessing the appropriateness of different core thresholds for uncovering stronger temporal signal, which is essential for subsequent phylodynamic analyses aiming to make epidemiological inference from time-stamped genome sequences (78, 79). Each dataset represents a different pathogen: *S. pneumoniae, N. gonorrhoeae, S.* Kentucky and *S. aureus* (10, 13, 17, 70). For each, we generated both pre-*Gubbins* (unfiltered) and post-*Gubbins* (recombination-filtered) alignments, which were processed with core thresholds ranging from 50% to 100% (see Methods).

In general, recombination-filtered alignments had fewer variant sites at each core threshold across all four datasets (Fig. 3a). Like the results observed in the large datasets, the number of variant sites for all recombination-filtered alignments were generally consistent until a decrease as the core threshold approached 100% (Supplementary Dataset 3). The unfiltered alignments had an overall higher number of variant sites, especially at lower core thresholds. Across all datasets, for both the recombination-filtered and unfiltered alignments, the lowest number of variant sites were observed in the 100% core threshold alignments. We identified specific instances in the unfiltered alignments, such as at 71% and 91% core thresholds for *S. aureus* and at 78% core for *S.* Kentucky, where a sharp decrease in variant sites occurred due to large regions of consistent sequence patterns across samples, rather than low-quality data. In each of these cases, the relevant genomic regions were masked by *Gubbins*, so these sharp decreases did not occur in the recombination-filtered alignments. Additionally, when comparing the tree topology generated from a 100% core threshold alignment to those from a 95% core threshold alignment, we observed some changes to the topology including more polytomous branching in the 100% core alignment, with other topological variation being minor (Supplementary Fig. 7-8, Supplementary Table 3). This suggests that the evolutionary relationships between genomes were less resolved at the 100% core threshold.

**Figure 3:**
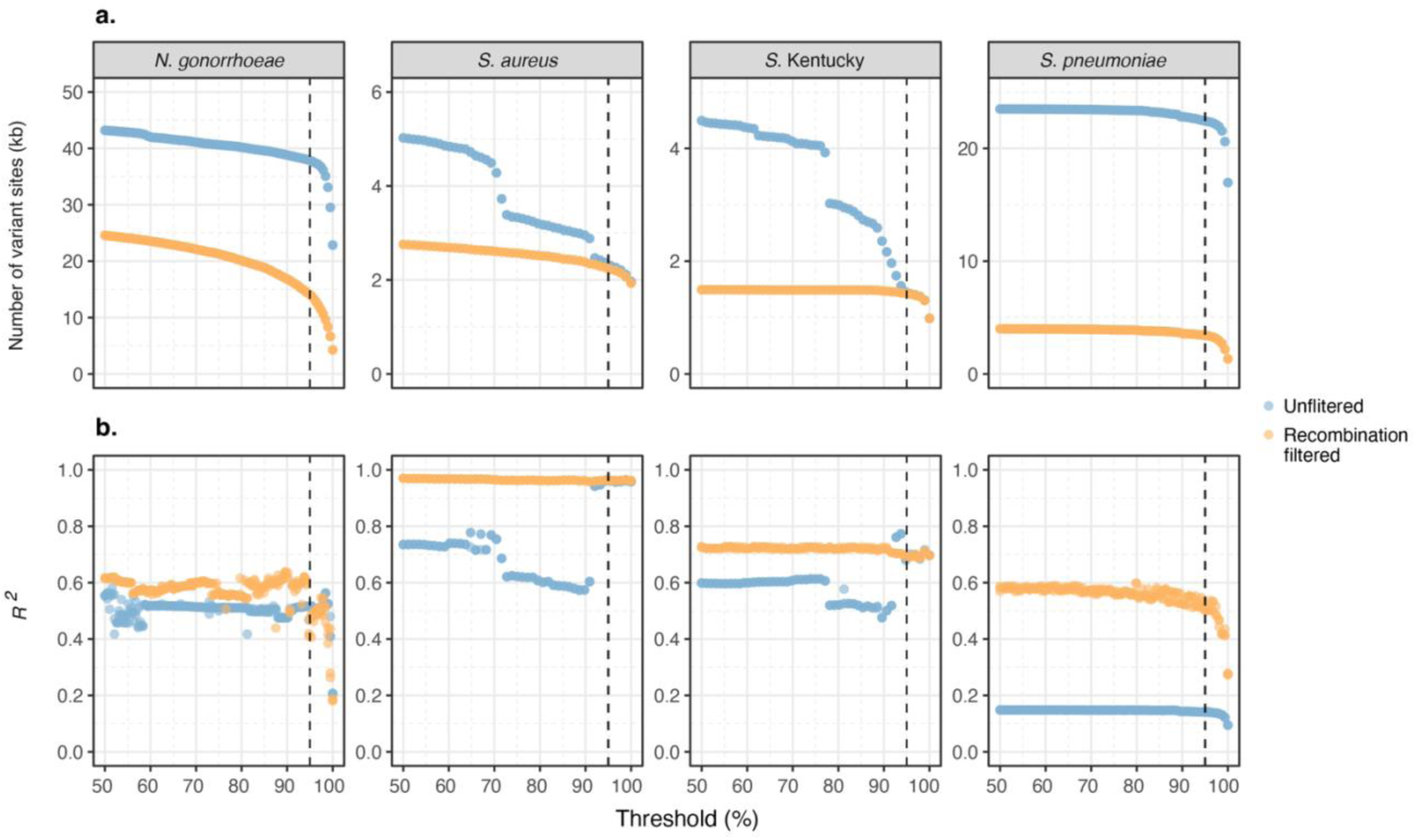
Impact of core thresholds on phylogenetic and temporal signal across bacterial datasets. (**a**) The number of variant sites (in kilobases) at each core genome threshold from 50% to 100% for four bacterial datasets: *Neisseria gonorrhoeae*, *Staphylococcus aureus*, *Salmonella enterica* subspecies Kentucky, and *Streptococcus pneumoniae*. The variant sites are shown for both recombination-filtered (using *Gubbins*) and unfiltered alignments. (**b**) The *R²* value from root-to-tip regression at each core threshold from 50% to 100% for phylogenetic trees constructed from the same four bacterial datasets, indicating the strength of the temporal signal. For all plots, the dotted line represents the 95% core threshold. Each dataset shows a steep decline in variant sites as the threshold approaches 100%, with *Neisseria gonorrhoeae* and *Streptococcus pneumoniae* also exhibiting a reduction in *R²* as the threshold approaches 100%.

Here, *R^2^* is a measurement of temporal signal in each dataset, which is essential for subsequent phylodynamic analyses aiming to estimate rates of epidemiological transmission. For the *N. gonorrhoeae* dataset, the *R*² values remained relatively stable across all tested core thresholds, until a steep decline as the core threshold approached 100%, with the same trend being observed for both the recombination-filtered and unfiltered phylogenies (Fig. 3b, Supplementary Dataset 3). Similarly, the *S. pneumoniae R*² values remained relatively stable across core thresholds, until a steep decline as the core threshold approached 100%. For the *S. pneumoniae* dataset, the impact of removing recombination on the *R*² values was more pronounced, with a mean *R*² difference of 0.41 between the recombination-filtered and unfiltered phylogenies. In the case of the *S. aureus* dataset, there was a noticeable difference between the *R*² values for the recombination-filtered and unfiltered phylogenies. Across all core thresholds, the recombination-filtered *S. aureus* phylogenies had very consistent *R*² values, while the *R*² values for the unfiltered *S. aureus* phylogenies were much more varied and did not follow a discernible trend. However, at the 100% core threshold, both the recombination-filtered and unfiltered *S. aureus* phylogenies had an *R*² value of 0.95. Similarly, the *S.* Kentucky dataset also showed relatively consistent *R*² values across all tested core thresholds for the recombination-filtered phylogenies and more varied *R^2^* values for the unfiltered phylogenies.

### Benchmarking Core-SNP-filter against existing tools

To evaluate the performance of *Core-SNP-filter* against three tools that perform related functions, *Goalign*, *SNP-sites* and *TrimAl*, we conducted benchmarking tests on two specific tasks: converting full alignments to no-invariant-sites alignments and converting no-invariant-sites alignments to 95% core alignments. *Core-SNP-filter* and *Goalign* can perform both tasks, whereas *SNP-sites* can only perform the first task, and *TrimAl* can only perform the second. These tools were chosen because they are written in fast, compiled programming languages: *Core-SNP-filter* in Rust, *Goalign* in Go, *SNP-sites* in C and *TrimAl* in C++.

There was a clear relationship between genome count and execution time for all tools and tasks (Fig. 4). For the task of removing invariant sites, *SNP-sites* was the fastest, with *Core-SNP-filter* taking about 1.5 times longer and *Goalign* taking about seven times longer (Fig. 4a). For the task of producing a 95% core alignment, *Core-SNP-filter* was the fastest, while *Goalign* and *TrimAl* took approximately four times longer (Fig 4c). In terms of RAM usage, *Core-SNP-filter* consistently used a fixed amount of RAM regardless of genome count for both removal of invariant sites (mean: 51.8 MB) and generating the 95% core alignments (mean: 2.8 MB) (Fig 4b and 4d). For removal of invariant sites, both *Goalign’s* and *SNP-sites’* RAM usage increased with genome count, reaching a mean of 78.8 GB and 709.1 MB, respectively, for the largest datasets (10,000 genomes). Similarly, for both *Goalign* and *TrimAl*, generating a 95% core threshold alignment resulted in increased RAM usage proportional to genome count, reaching a mean of 4.2 GB and 2.3 GB, respectively, for the largest datasets. Goalign loads the whole alignment into memory, which allows it to be used in a Unix pipe (accept input via stdin) but increases RAM usage. The other tools cannot operate in a pipe as they require multiple passes over the file.

**Figure 4:**
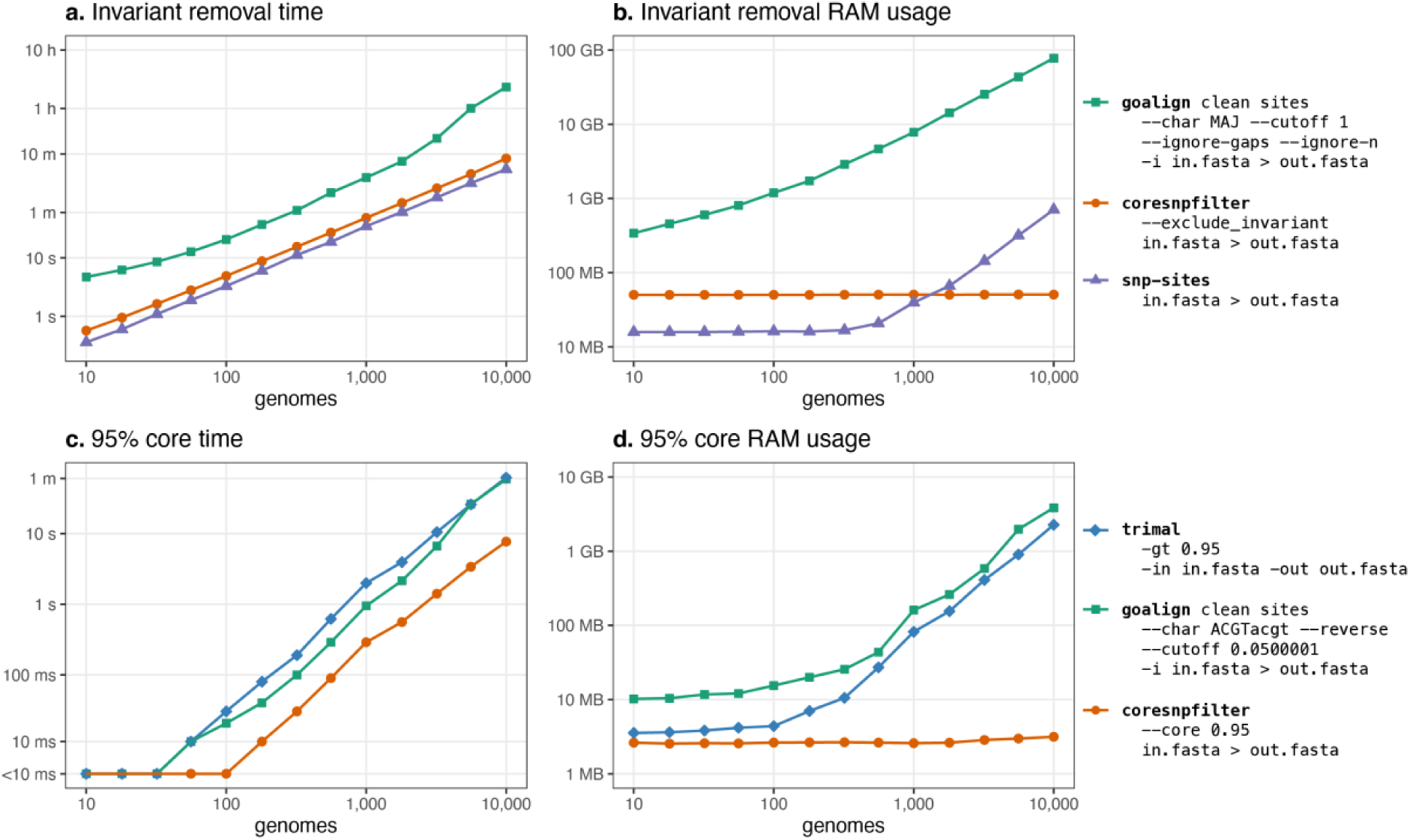
Benchmarking of Core-SNP-filter against other tools with similar functionality. The time taken (**a**) and RAM usage (**b**) to remove invariant sites from alignments with varying numbers of genomes using three different tools (*Goalign*, *Core-SNP-filter* and *SNP-sites*). The time taken (**c**) and RAM usage (**d**) to generate a 95% core alignment across alignments with varying numbers of genomes using three different tools (*TrimAl*, *Goalign and Core-SNP-filter*). All axes are transformed with a log scale.

## DISCUSSION

This study highlights the importance of considering soft-core strategies as a best practice for phylogenetic analyses of bacterial datasets. Our findings reveal the limitations of using a strict-(100%) core threshold, particularly for large and diverse datasets, where even a slight relaxation from a 100% core to 99% notably increases the number of output sites. A 95% core emerges as a practical choice for many scenarios, striking a balance between retaining informative phylogenetic signals and excluding data inconsistent with the clonal frame of evolution or influenced by recombination. We also benchmarked the computational performance of our tool, *Core-SNP-filter*, against available tools that perform part, or all, of the required processing, demonstrating its efficiency in handling large datasets.

Traditional methods often rely on generating strict alignments from all genomes in a dataset as the basis for phylogenetic inferences and genomics evolutionary studies. However, as we demonstrate, this approach becomes increasingly impractical and ineffective as datasets grow. The strict-core method, which requires alignment of SNPs present in all genomes, often results in significant data loss, reducing the resolution and accuracy of phylogenetic analyses. Recent studies have explored alternative methods for generating phylogenies from large bacterial datasets, including concatenated core genome multilocus sequence typing (cgMLST) alignments and neighbour-joining trees based on pairwise core SNP distances (19, 80, 81). While these methods address some challenges, our study presents a simple tool striking a balance between traditional strict-core approaches and addressing present limitations. These soft-core approaches are being implemented in a number of large-scale phylogenetic studies (6, 26, 27).

In public health and hospital outbreak settings, phylogenetic analysis is crucial for inferring transmission dynamics. Typically, SNP distances calculated from strict-core alignments are used to determine transmission clusters. However, this method can be problematic due to its sensitivity to reference genome selection, masking of recombinogenic or prophage regions, and the approach taken (23). These issues are exacerbated in large datasets, where a static SNP distance threshold for transmission clustering may be inappropriate. Such thresholds, while standard in many transmission studies and accredited pipelines, fail to account for dynamic core sizes that result from strict-core approaches, particularly in epidemiological studies involving expanding datasets (82–84). Our findings indicate that as datasets expand, the alignment size diminishes under a strict-core threshold, potentially leading to over-supported transmission networks and misleading clinical or public health conclusions. Moreover, the use of a strict-core approach resulted in a higher proportion of identical genome pairs across multiple studies, suggesting a loss of resolution that could obscure important genetic distinctions. This reduced resolution may impact decisions in outbreak response, where the ability to detect subtle genetic differences is essential for accurate transmission inference. By relaxing the core threshold, stable alignment sizes can be maintained up to and likely beyond 10,000 genomes, improving phylogenetic accuracy without compromising data integrity.

Furthermore, the strict-core approach poses challenges when generating whole-genome alignments from enrichment or capture-based methods, which are valuable for sequencing unculturable pathogens but often result in higher rates of missing data. These enrichment and capture-based methods are relatively new and are being used more frequently for bacteria, necessitating the establishment of best-practice data analysis techniques (6, 11, 85, 86). Similarly, missing data may arise when sequencing viral genomes, especially when genomes are sequenced directly from clinical samples. The inherent diversity of viruses and often degraded nature of clinical samples exacerbate the problem, making it more challenging to achieve complete sequencing coverage (87). As a result, adjustments to account for missing data have become more prevalent in viral and enrichment WGS data, driving the adoption of more adaptable analytical methods. For these datasets, a soft-core approach conserves data that are part of the clonal frame of evolution but sometimes fail to sequence (88). To address this issue, enrichment and viral studies have started to adopt soft-core methods using alternative tools, highlighting the need for more flexible approaches to preserve critical genomic information (6, 11).

For datasets with a strong temporal signal and limited geographic diversity, discarding SNP positions using a strict-core threshold can remove information used by phylodynamic analyses. A soft-core approach can be valuable here and complement other tools such as *Gubbins* for recombination filtering. Our analysis showed that for recombination-unfiltered alignments, a 95% core threshold generally performs better (e.g., *S. pneumoniae* and *N. gonorrhoeae*) or similarly (e.g., *S. aureus* and *S.* Kentucky) to a 100% core threshold, making 95% the preferred threshold. Lower thresholds sometimes performed worse on unfiltered alignments due to recombinant regions. In our *S. aureus* dataset, for instance, one isolate (SAMEA3212906) appeared as an outlier on a long branch in the root-to-tip at lower thresholds (<92%), whereas higher thresholds (>92%) helped to isolate the phylogenetic signal. For recombination-filtered alignments, the core threshold showed little impact on temporal signal values, except for sometimes causing a clear decline near 100%. Thresholds ranging from 50% to 95% performed similarly, suggesting that overly strict cores discard valuable data. For example, our recombination-filtered *N. gonorrhoeae* dataset lost over 90% of SNPs at a 100% core threshold. Overall, a 95% core threshold generally outperforms a 100% threshold.

The impact of missing data on phylogenetic inference has been a topic of significant ongoing debate, with studies showing contrasting views (24, 25, 89, 90). Several studies find that minimal missing data does not substantially affect phylogenetic outcomes, especially in large datasets where missing data is proportionally low (25, 89). However, other studies indicate that higher levels of missing data, such as >50% of sites, can introduce uncertainty in inferred topologies and result in misleading branch lengths (24, 90). A further consideration is the datasets used to explore this question, which range from short alignments of *18S* and *28S* rRNA from tapeworms to long alignments in the thousands for brush-tongued parrots (24, 25). Overall, these studies debate the utility of including some missing data in datasets of differing scale and origin. Here, we instead focus on the fundamental point that including sites with some gaps provides additional phylogenetic information for those samples where the site is known. We show that phylogenetic trees constructed from a 95% core threshold alignment can result in better-resolved topologies than those from 100% core threshold alignments. In the context of Felsenstein’s pruning algorithm, which underpins likelihood based phylogenetics, sites with some gaps carry information to resolve relationships between samples for which a base is present that would otherwise be discarded under a 100% core threshold, while gaps do not reduce the information offered by these terms (91). Depending on the data at hand, additional sites with some gaps may serve to make estimated branch lengths and topology more or less certain, depending on likelihoods for a branch length or a given topology at sites including gaps. Thus, our approach of using a 95% core threshold can result in better-resolved topologies, particularly in large-scale bacterial datasets.

When comparing *Core-SNP-filter* to other tools with similar capabilities, the choice of tool has minimal impact for small datasets (e.g., <100 samples) as all tested tools are sufficiently fast and efficient. However, for large datasets (e.g., >1000 samples), especially those containing invariant sites, both time and RAM become critical factors, and *Core-SNP-filter* has the potential to scale to very large datasets (e.g., >100,000 samples). *Goalign* presents challenges due to its non-intuitive commands for excluding invariant sites and producing soft-core alignments, though it has functionality for processing alignments in ways that *Core-SNP-filter* cannot.

Our study underscores the advantages of using soft-core thresholds, particularly the 95% threshold, over 100% strict-core thresholds for bacterial genome datasets. The soft-core approach, achieved with *Core-SNP-filter,* preserves more informative sites, enhancing phylogenetic resolution and temporal signal, while being computationally efficient. Researchers and public health professionals should consider using a soft core in their analyses to retain available evolutionary signal and accurately interpret phylogenetic relationships and transmission patterns.

## ETHICAL APPROVAL

No ethics was required for this study.

## AUTHOR CONTRIBUTIONS

MLT, RRW, GT, and LAF designed the study. MLT and RRW performed all analyses. MLT, RRW, TS, GT and LAF prepared the figures. GT, DJI, TS and TPS provided supervision and support. The first draft of the manuscript was written by MLT, being revised by all authors. All authors had full access to all study data. The corresponding authors had final responsibility for the decision to submit for publication.

