## Supplementary Appendix for "Exploring SNP Filtering Strategies: The Influence of Strict vs Soft Core"

**Supplementary Figure 1**

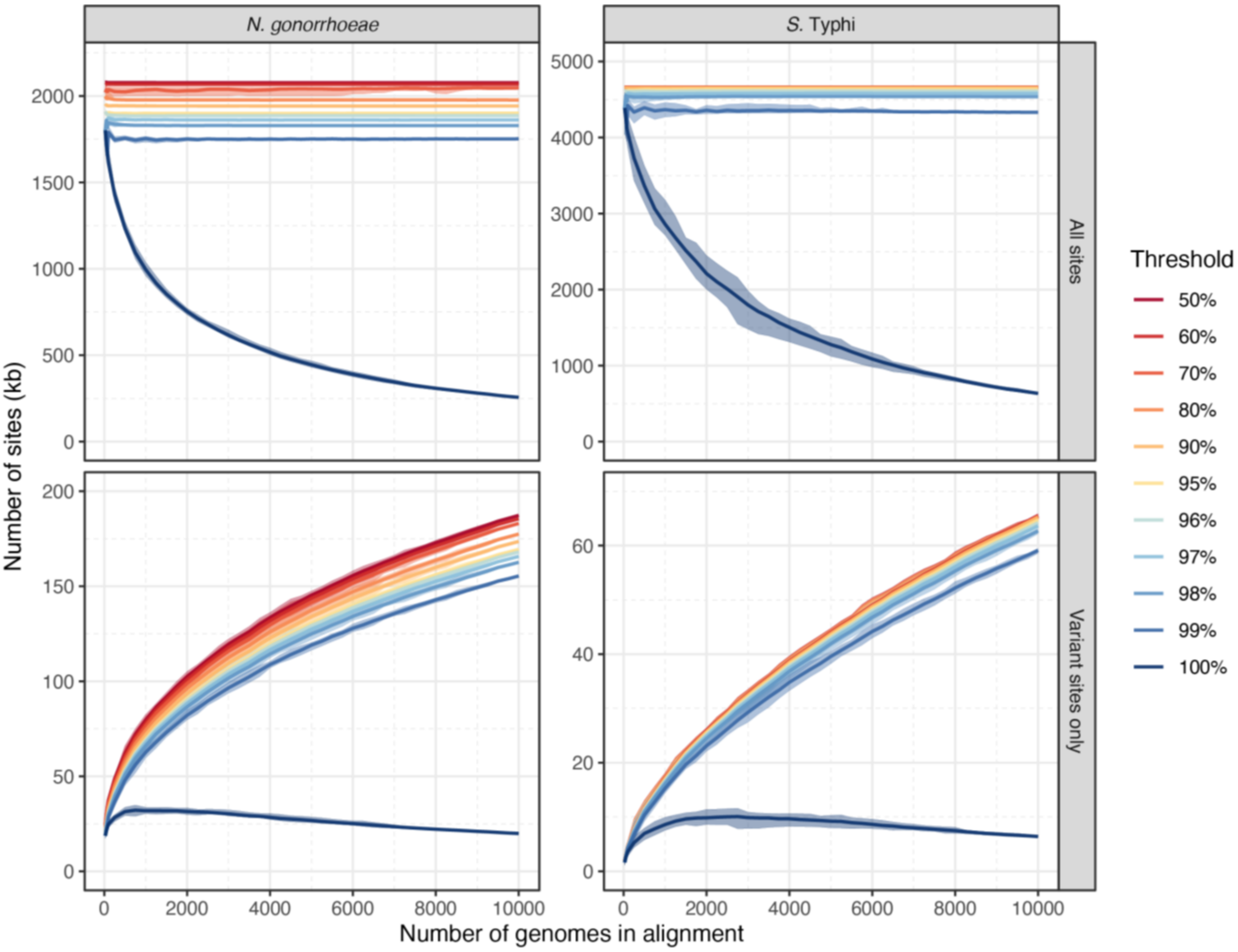


The number of alignment sites (in kilobases) is plotted against the number of genomes included in each alignment (ranging from 25 to 10,000). Each alignment was processed at core thresholds ranging from 50% to 100% (indicated by line colour). All sites in *N. gonorrhoeae* alignments, all sites in *S.* Typhi alignments, variant sites only in *N. gonorrhoeae* alignments, and variant sites only in *S.* Typhi alignments. The mean of ten alignments consisting of randomly aggregated genomes is plotted as the solid line and the minimum to maximum range for each threshold is plotted in the shaded area around each line.

**Supplementary Figure 2**

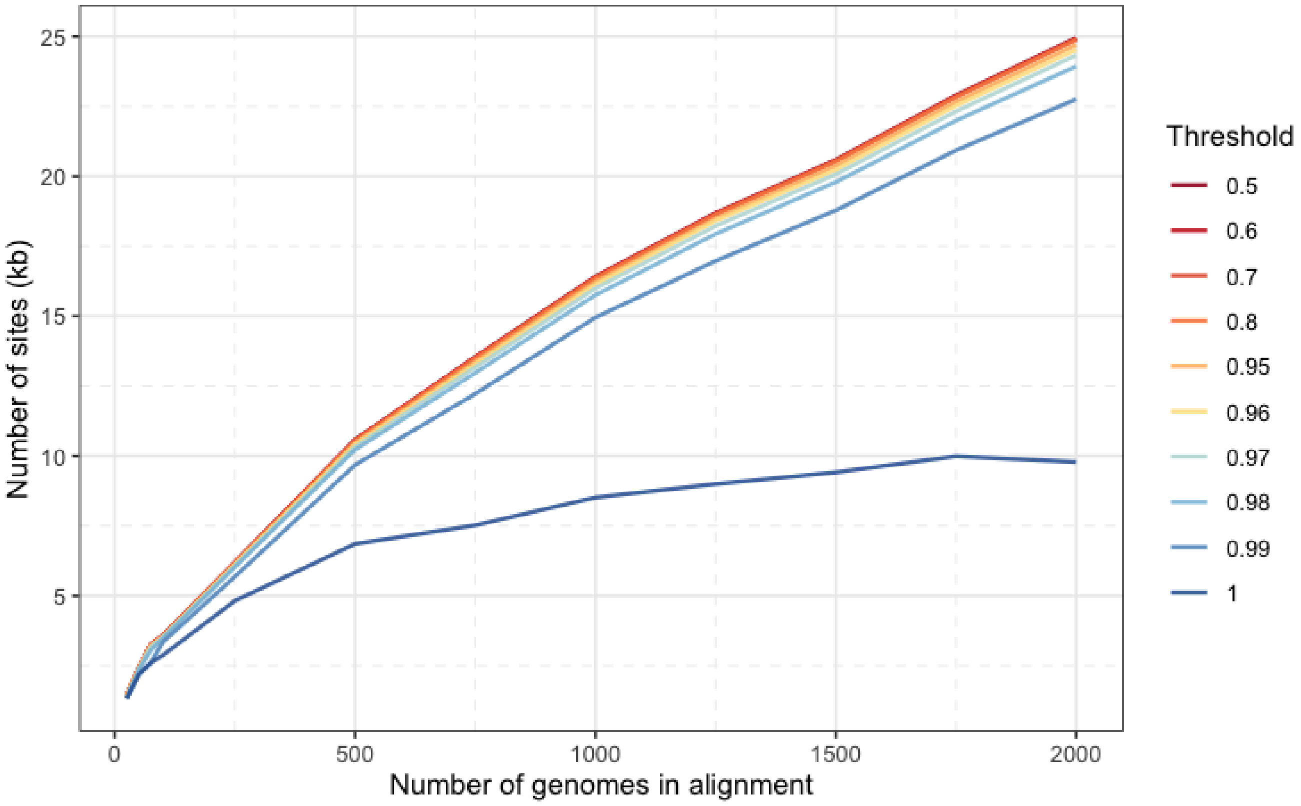


The number of variant sites in *S.* Typhi alignments with various numbers of genomes. Following removal of invariant and recombinogenic regions, each alignment was processed at core thresholds ranging from 50% to 100% (indicated by line colour) and the number of variant sites plotted. *S.* Typhi was chosen as recombination detection works best on closely related genomes, and for computational efficiency, only one set of alignments was only tested up until the 2000-genome alignment.

**Supplementary Figure 3**

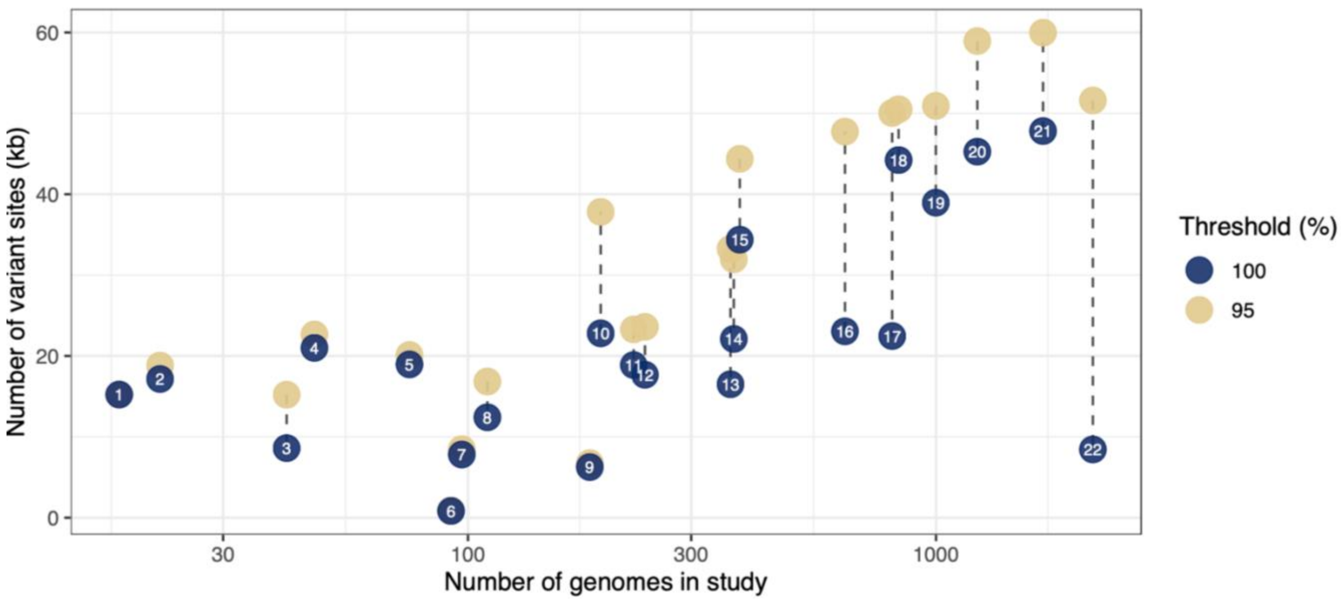


The number of variant sites (in kilobases) for each *N. gonorrhoeae* dataset tested at both a 100% and 95% core threshold alignment. The labels refer to datasets as follows: 1. Ezewudo *et al.* 2015, 2. Wind *et al.* 2017, 3. Ryan *et al.* 2018, 4. Kwong *et al.* 2016, 5. Kwong *et al.* 2018 6. Buckley *et al.* 2018, 7. Fifer *et al.* 2018, 8. Cehovin *et al.* 2018, 9. Didelot *et al.* 2016, 10. Golparian *et al.* 2020, 11. Lan *et al.* 2020, 12. Yahara *et al.* 2018, 13. Demczuk *et al.* 2015, 14. Lee *et al.* 2018, 15. Sanchez-Buso *et al.* 2019, 16. Thomas *et al.* 2019, 17. Alfsnes *et al.* 2020, 18. Mortimer *et al.* 2020, 19. Grad *et al.* 2016, 20. Town *et al.* 2020, 21. DeSilva *et al.* 2016, 22. Williamson *et al.* 2019.

**Supplementary Figure 4**

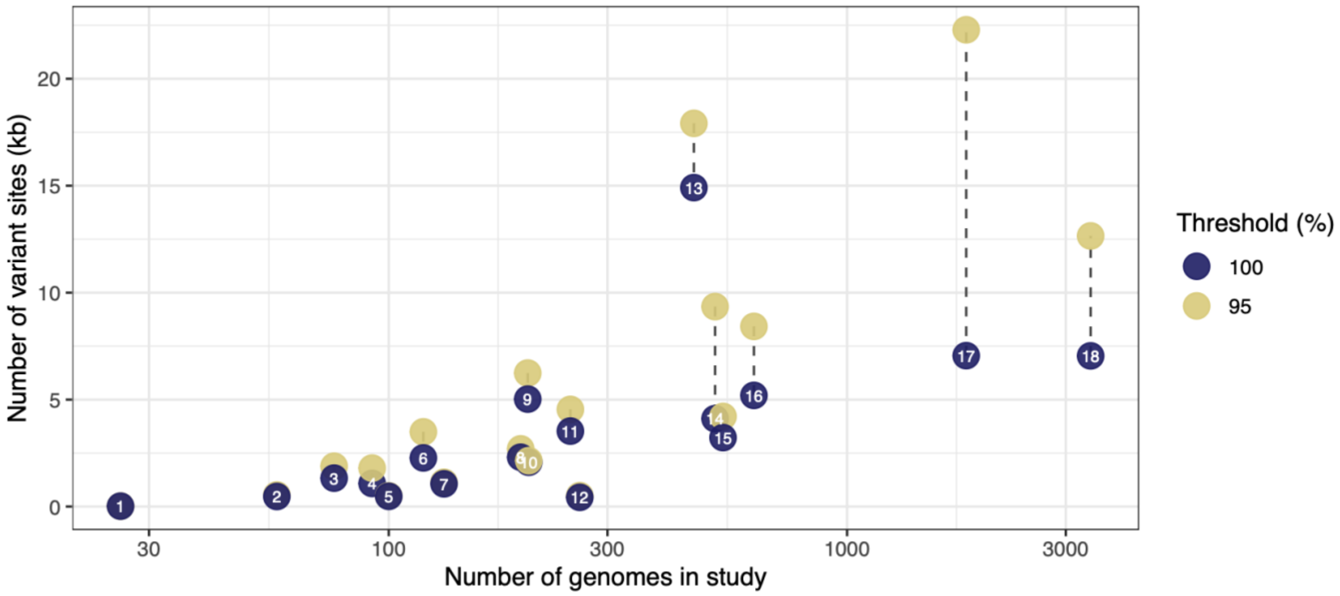


The number of variant sites (in kilobases) for each *S*. Typhi dataset tested at both a 100% and 95% core threshold alignment. The labels refer to datasets as follows: 1. Rasheed *et al*. 2020, 2. Thilliez *et al*. 2022, 3. Guevara *et al*. 2021, 4. Argimon *et al*. 2022, 5. Klemm *et al*. 2018, 6. Ingle *et al*. 2019, 7. Karuiki *et al*. 2021, 8. Pragasam *et al*. 2020, 9. Maes *et al*. 2022, 10. Rahman *et al*. 2020, 11. Park *et al*. 2018, 12. Gauld *et al*. 2022, 13. Carey *et al*. 2024, 14. Ashton *et al*. 2016, 15. Tanmoy *et al*. 2018, 16. Chattaway *et. al* 2021, 17. Wong *et al*. 2015, 18. Da Silva *et al*. 2022.

### Supplementary Figure 5

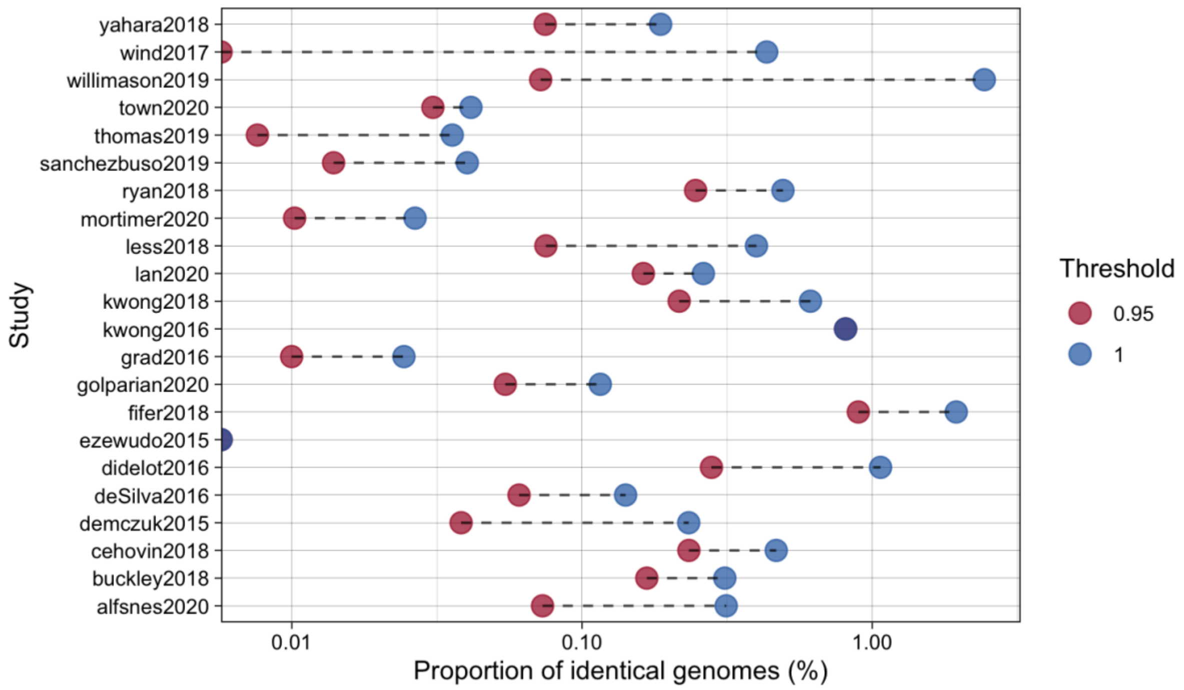


The plot shows the proportion of genome pairs with a pairwise SNP distance of zero for each *N. gonorrhoeae* dataset, calculated as a percentage of all possible genome pairs. Proportions are displayed for alignments processed at two core SNP thresholds: 100% and 95%. Dashed lines connect the values for the two thresholds within the same dataset, illustrating differences between the thresholds. The x-axis is presented on a logarithmic scale to highlight variations across datasets with differing proportions.

**Supplementary Figure 6**

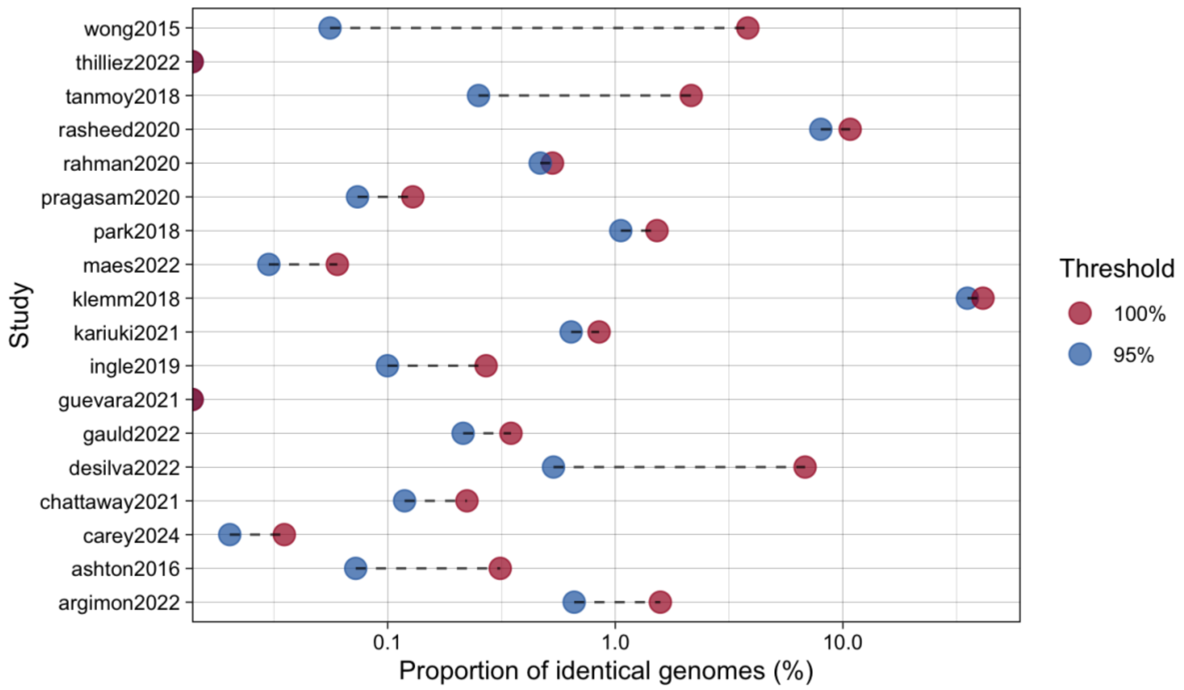


The plot shows the proportion of genome pairs with a pairwise SNP distance of zero for each *S*. Typhi dataset, calculated as a percentage of all possible genome pairs. Proportions are displayed for alignments processed at two core SNP thresholds: 100% and 95%. Dashed lines connect the values for the two thresholds within the same dataset, illustrating differences between the thresholds. The x-axis is presented on a logarithmic scale to highlight variations across datasets with differing proportions.

### Supplementary Figure 7

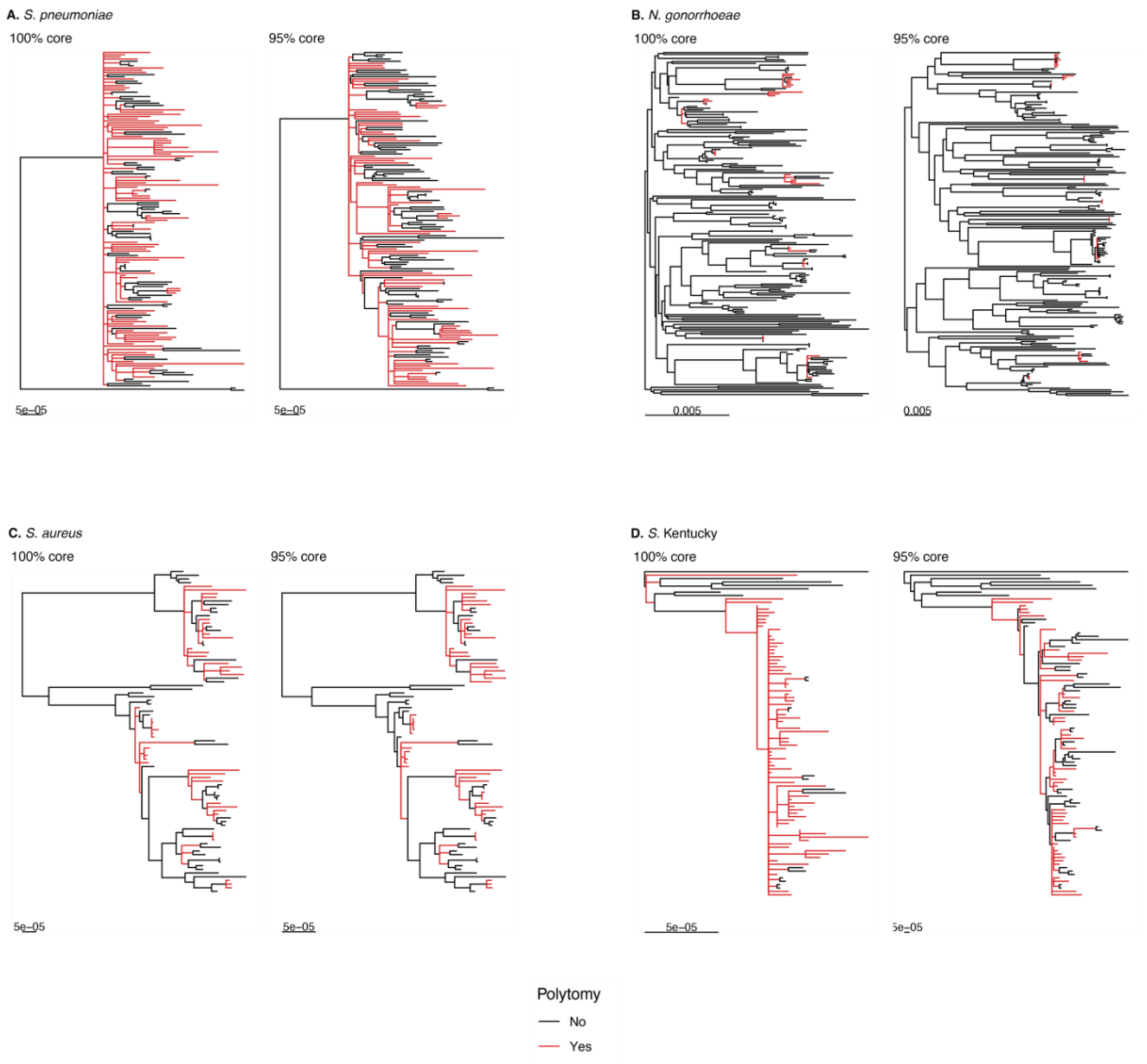


Midpoint rooted phylogenetic trees for the four datasets used in the temporal analysis of this study. For each dataset a phylogenetic tree resulting from both the 100% and 95% core threshold alignments is presented. Phylogenetic trees were generated allowing for polytomies, and these branches are coloured in red for all trees. The number of polytomies in each phylogeny can be found in Supplementary Table 3.

### Supplementary Figure 8

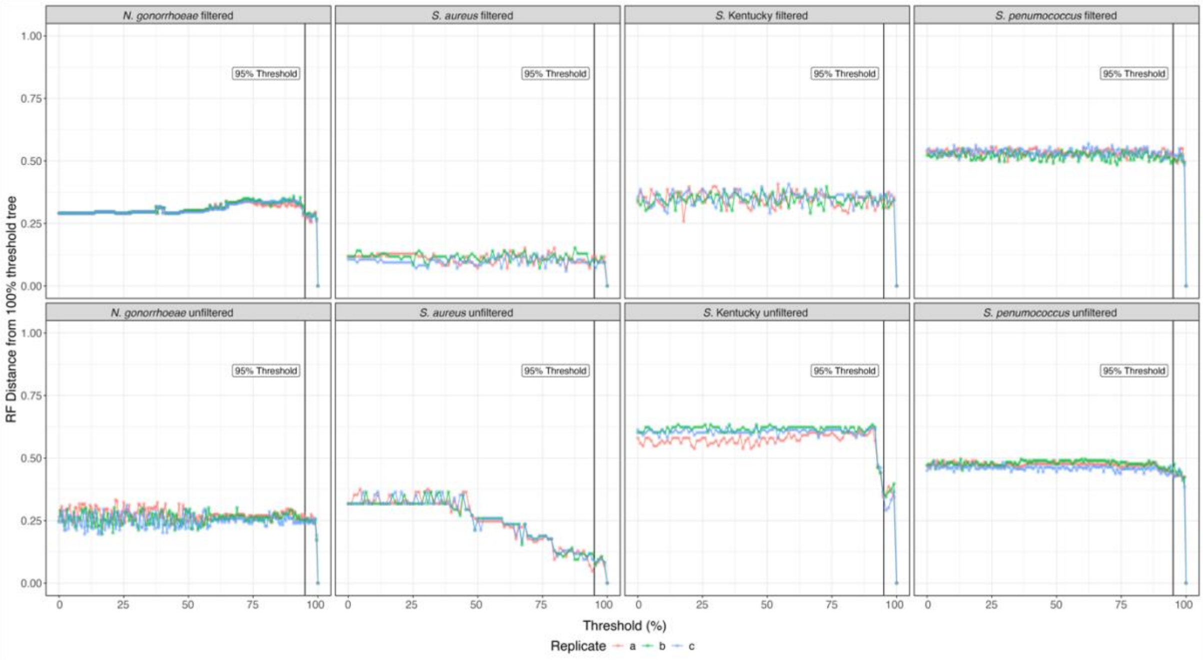


Robinson-Foulds (RF) distances compare the topologies of phylogenies generated using a 100% strict-core threshold to those generated at varying thresholds (0–99%) across three replicates per dataset. RF distances are normalized: a value of one indicates maximally dissimilar trees, while zero indicates identical topologies. "Filtered" phylogenies were based on recombination-filtered alignments, while "unfiltered" phylogenies used unfiltered alignments. Across datasets, RF distances remain low as the core threshold decreases from 100%, suggesting that non-core information enhances the phylogenetic signal rather than distorting it.

**Supplementary Table 1: *N. gonorrhoeae* study alignments using 100% and 95% core-thresholds**

| **Dataset** | **Variant sites at 100% core threshold** | **Variant sites at 95% core threshold** | **Percentage identical pairs at 100% core threshold (%)** | **Percentage identical pairs at 95% core threshold (%)** | **Number of genomes in study** | **Reference** |
|---|---|---|---|---|---|---|
| Alfsnes *et al.* 2020 | 22465 | 50042 | 0.31 | 0.07 | 806 | [1] |
| Buckley *et al.* 2018 | 828 | 905 | 0.31 | 0.17 | 92 | [2] |
| Cehovin *et al.* 2018 | 12414 | 16838 | 0.47 | 0.23 | 110 | [3] |
| Demczuk *et al.* 2015 | 16500 | 33240 | 0.23 | 0.04 | 364 | [4] |
| DeSilva *et al.* 2016 | 47813 | 59973 | 0.14 | 0.06 | 1692 | [5] |
| Didelot *et al.* 2016 | 6257 | 6747 | 1.07 | 0.28 | 182 | [6] |
| Ezewudo *et al.* 2015 | 15225 | 15225 | 0.00 | 0.00 | 18 | [7] |
| Fifer *et al.* 2018 | 7812 | 8483 | 1.95 | 0.90 | 97 | [8] |
| Golparian *et al.* 2020 | 22798 | 37817 | 0.12 | 0.05 | 192 | [9] |
| Grad *et al.* 2016 | 38953 | 50924 | 0.02 | 0.01 | 999 | [10] |
| Kwong *et al.* 2016 | 20994 | 22655 | 0.81 | 0.81 | 47 | [11] |
| Kwong *et al.* 2018 | 18954 | 20122 | 0.61 | 0.22 | 75 | [12] |
| Lan *et al.* 2020 | 18879 | 23281 | 0.26 | 0.16 | 226 | [13] |
| Lee *et al.* 2018 | 22100 | 31954 | 0.40 | 0.08 | 370 | [14] |
| Mortimer *et al.* 2021 | 44221 | 50502 | 0.03 | 0.01 | 832 | [15] |
| Ryan *et al.* 2018 | 8600 | 15209 | 0.49 | 0.25 | 41 | [16] |
| Sanchez-Buso *et al.* 2019 | 34379 | 44371 | 0.04 | 0.01 | 381 | [17] |
| Thomas *et al.* 2019 | 23045 | 47741 | 0.04 | 0.01 | 639 | [18] |
| Town *et al.* 2020 | 45262 | 58937 | 0.04 | 0.03 | 1225 | [19] |
| Willimason *et al.* 2019 | 8441 | 51593 | 2.43 | 0.07 | 2162 | [20] |
| Wind *et al.* 2017 | 17148 | 18816 | 0.43 | 0.00 | 22 | [21] |
| Yahara *et al.* 2018 | 17676 | 23605 | 0.19 | 0.07 | 239 | [22] |

**Supplementary Table 2: *S.* Typhi study alignments using 100% and 95% core-thresholds**

| Dataset | Variant sites at 100% core threshold | Variant sites at 95% core threshold | Percentage identical pairs at 100% core threshold (%) | Percentage identical pairs at 95% core threshold (%) | Number of genomes in study | Reference |
|---|---|---|---|---|---|---|
| Argimon *et al.* 2022 | 1080 | 1798 | 1.58 | 0.66 | 92 | [23] |
| Ashton *et al.* 2016 | 4110 | 9349 | 0.31 | 0.07 | 515 | [24] |
| Carey *et al.* 2024 | 14911 | 17915 | 0.04 | 0.02 | 463 | [25] |
| Chattaway *et. al* 2021 | 5197 | 8423 | 0.22 | 0.12 | 626 | [26] |
| Da Silva *et al.* 2022 | 7045 | 12652 | 6.83 | 0.54 | 3398 | [27] |
| Gauld *et al.* 2022 | 430 | 509 | 0.35 | 0.21 | 261 | [28] |
| Guevara *et al.* 2021 | 1328 | 1886 | 0.00 | 0.00 | 76 | [29] |
| Ingle *et al.* 2019 | 2274 | 3493 | 0.27 | 0.10 | 119 | [30] |
| Karuiki *et al.* 2021 | 1061 | 1134 | 0.85 | 0.64 | 132 | [31] |
| Klemm *et al.* 2018 | 478 | 501 | 41.29 | 35.25 | 100 | [32] |
| Maes *et al.* 2022 | 5023 | 6241 | 0.06 | 0.03 | 201 | [33] |
| Park *et al.* 2018 | 3528 | 4543 | 1.53 | 1.06 | 249 | [34] |
| Pragasam *et al.* 2020 | 2306 | 2697 | 0.13 | 0.07 | 194 | [35] |
| Rahman *et al.* 2020 | 2071 | 2145 | 0.53 | 0.47 | 202 | [36] |
| Rasheed *et al.* 2020 | 19 | 20 | 10.77 | 8.00 | 26 | [37] |
| Tanmoy *et al.* 2018 | 3223 | 4214 | 2.16 | 0.25 | 536 | [38] |
| Thilliez *et al.* 2022 | 470 | 540 | 0.00 | 0.00 | 57 | [39] |
| Wong *et al.* 2015 | 7055 | 22289 | 3.82 | 0.06 | 1819 | [40] |

**Supplementary Table 3: Number of polytomies in phylogenies generated from alignments at a 100% core SNP threshold compared to those generated from alignments at a 95% core SNP threshold for four bacterial datasets.**

| **Dataset** | **Core SNP threshold** | **Number of nodes** | **Number of polytomies** | **Percentage (%)** |
|---|---|---|---|---|
| *S. pneumoniae* | 100% | 226 | 127 | 56.19 |
| *S. pneumoniae* | 95% | 248 | 109 | 43.95 |
| *N. gonorrhoeae* | 100% | 365 | 40 | 10.96 |
| *N. gonorrhoeae* | 95% | 369 | 36 | 9.76 |
| *S. aureus* | 100% | 148 | 63 | 42.57 |
| *S. aureus* | 95% | 146 | 67 | 45.89 |
| *S.* Kentucky | 100% | 121 | 94 | 77.69 |
| *S.* Kentucky | 95% | 157 | 70 | 44.59 |
